# Appia: simpler chromatography analysis and visualization

**DOI:** 10.1101/2022.04.01.486784

**Authors:** Richard Posert, Isabelle Baconguis

## Abstract

Chromatography is an essential family of assays for molecular biology and chemistry. Typically, only a qualitative assessment of peak height, position, and shape are sufficient to proceed. However, there are more advanced forms of chromatography that require integration of peak width, baseline subtraction, and precise peak position calling. To handle these precise workflows, instrument manufacturer software is complicated and arcane; this makes simpler day-to-day analyses inefficient. Additionally, each manufacturer reports the data in their own proprietary format, requiring that analysis is performed at instrument computers and making data sharing between labs difficult. Here we present Appia, a free, open-source chromatography processing and visualization package focused on making analysis, collaboration, and publication quick and easy.

## Introduction

Chromatography is a foundational assay for investigating the quantity, homogeneity, and purity of small molecule and protein samples. In its most basic form, chromatography involves measuring the spatial separation of an analyte induced by flowing a mobile phase over a stationary matrix (1). While the mobile phase flows through the matrix, sample is carried along at a varying rate based on the property to be analyzed. For example, in size-exclusion chromatography (SEC), the smaller analytes move more slowly through the packed resin than larger ones. Thus, peak position (that is, the time or volume at which sample elutes) informs the user about the relative size of the analyte, but in reverse-phase chromatography peak position correlates with the relative hydrophilicity of the analyte (2,3). Peak shape gives information about the homogeneity of the species with regard to the measured property. Peak area gives the total amount of analyte injected, but peak height is often used as a heuristic for peak area (which can be difficult to accurately measure). For most experiments, the peak position, height, and width information about the analyte sufficient for proceeding with the next experiment.

The simplicity of these three main readouts means that most day-to-day chromatography experiments can be satisfactorily analyzed by eye in a few seconds. Indeed, screening data from a type of SEC ubiquitous in our field of structural biochemistry are often laid out in a large grid of very small plots because sample quality is so immediately recognizable (4). This layout hints at an issue --- simple assays require simple presentation to be legible. Software written by chromatogram manufacturers must be capable of the less-common, complex chromatographic assays and so is in conflict with a simple presentation of the data. Moreover, each chromatogram manufacturer develops their own analysis software, meaning that data formats are not inter-operable, and often users must switch between two or more different programs to analyze a single experiment. This appears a simple inconvenience, but there is a great body of research indicating that interface complexity and “task-switching” (in this case, switching between two different manufacturer software packages) wastes effort and makes learning and insight more difficult (5-7). Given how common chromatography is, even small wastes of time or energy add up to a significant delay over the length of a project. We wrote Appia to solve these problems by giving simple assays a simple interface.

## Results

Appia is a free, open-source chromatography package which converts data from all major chromatography instruments (Waters, Agilent, Shimadzu, and GE/Cytiva) into a standard, “tidy” format (8). First, we will describe the actual functionality that Appia provides. Next, we will demonstrate its utility in day-to-day chromatography experiments, with a focus on protein biochemistry. We focus on protein biochemistry because that is the main focus of our lab. However, Appia can perform the same process on any supported chromatogram’s data.

Before they can be visualized, chromatography data must be processed into an Appia “Experiment”. The user provides raw data files exported from their chromatography instrument. Appia will prompt the user for some basic information (flow rate, detection channel, etc.) about the run, if that information is not included in the data files. Appia then processes this data into a standard “tidy” data format which is written to a CSV file. After data are processed, they can be visualized. The user can perform their own visualizations for more advanced experiments using the CSV file, but Appia provides functionality to automatically create appealing basic plots during processing. Additionally, if users set up an Appia Web server, the data will be uploaded to the Appia Web database for viewing through a browser.

The Appia Web interface provides a centralized location for viewing all experiments processed with Appia (Supplementary Video). A demo of Appia Web is available at http://traces.baconguislab.com. Users can scroll through an experiment list, or type to search. Once selected, the data are displayed in zoomable, interactive line plots. Data are separated into analytic (Figure 2A) and preparative (Figure 2B) data, since these experiments typically require different visualization. For preparative data, the overall trace is displayed with fractions highlighted in color below it. Analytic data is displayed with a single line per injection, colored by injection name. Appia can handle any number of simultaneous acquisition channels, which are all plotted separately. Users can click the color legend to turn on and off display of fractions or traces. Additionally, users can zoom in to a region of the trace and normalize the maximum value of each injection in that region to 1. These normalized plots, presented alongside the raw-data plots, make it easy to compare peaks from different injections to a reference peak. Once a satisfactory image is created, it can be downloaded directly from the web interface.

**Figure 1.**
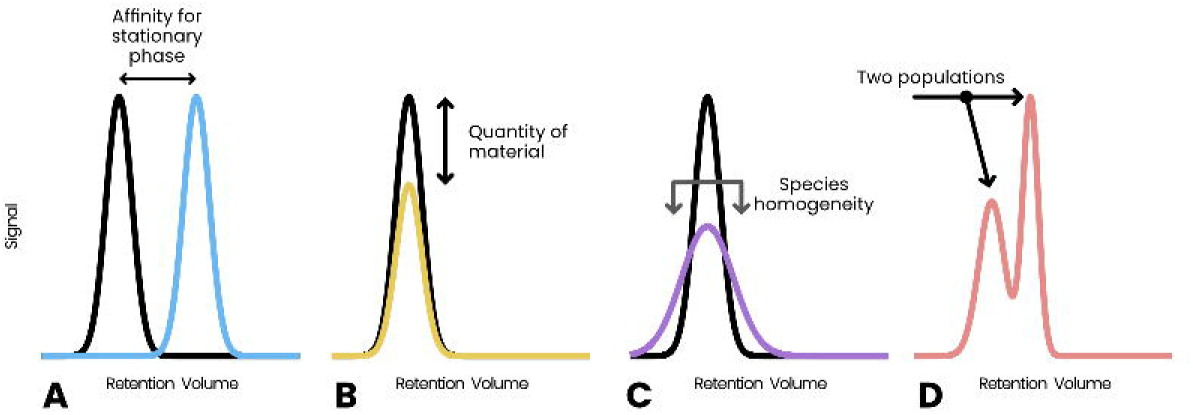
The three main properties of gaussian chromatography peaks. Peak position, or retention volume (A) informs of the relative affinity for the mobile and stationary phases. The peak height (B) informs of the amount of material included in the injection. The peak shape (C) informs of the heterogeneity of the species. If two peaks (D) typically represents two distinct species

**Figure 2.**
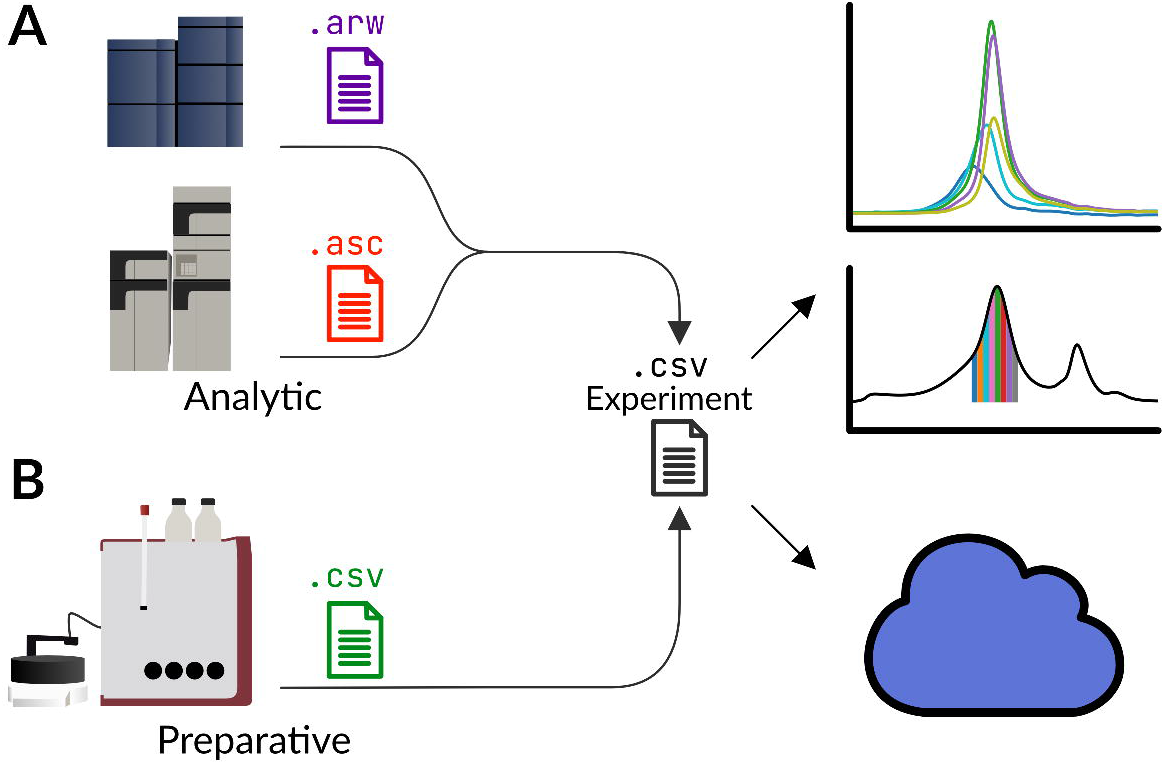
Appia processes proprietary data formats (colored) from a variety of manufacturers and automatically determines whether the data are analytic (A) or preparative (B). Appia saves this data in standardized CSV format files (black) for plotting. Chromatography data are uploaded to Appia Web at the same time.

If a user selects multiple experiments, they are combined. For analytic data, injections are labeled with their respective experiment names and plotted together for ease of comparison. Preparative data for combined experiments show the overall injection profile colored by experiment without fraction fills. As with single experiment display, all graphs are fully interactive. The list of selected experiments, X-coordinate zoom, and normalization region are continuously stored and updated in the URL query string. This means that a link shares not only the list of experiments, but a particular region of the trace to be inspected (for example: http://traces.baconguislab.com/HPLC_Example_1+HPLC_Example_2?view-range=1.93-2.13).

All of the above make chromatography data easier to work with, but thinking of it as merely a tool of convenience ignores difficulty of gaining insight from data which are difficult to work with. For instance, a common assay in our lab is exploration of expression levels of a given protein construct. We infect cells with a virus, causing them to express a fluorescently-tagged protein, which we track using SEC. The ideal expression level is often a function of MOI, additive concentration, expression time, and temperature. Appia’s standard data format means that users can write a processing script or develop a Prism/Excel workflow once, and then apply it quickly to the data every time this assay is run (Figure 3A).

**Figure 3.**
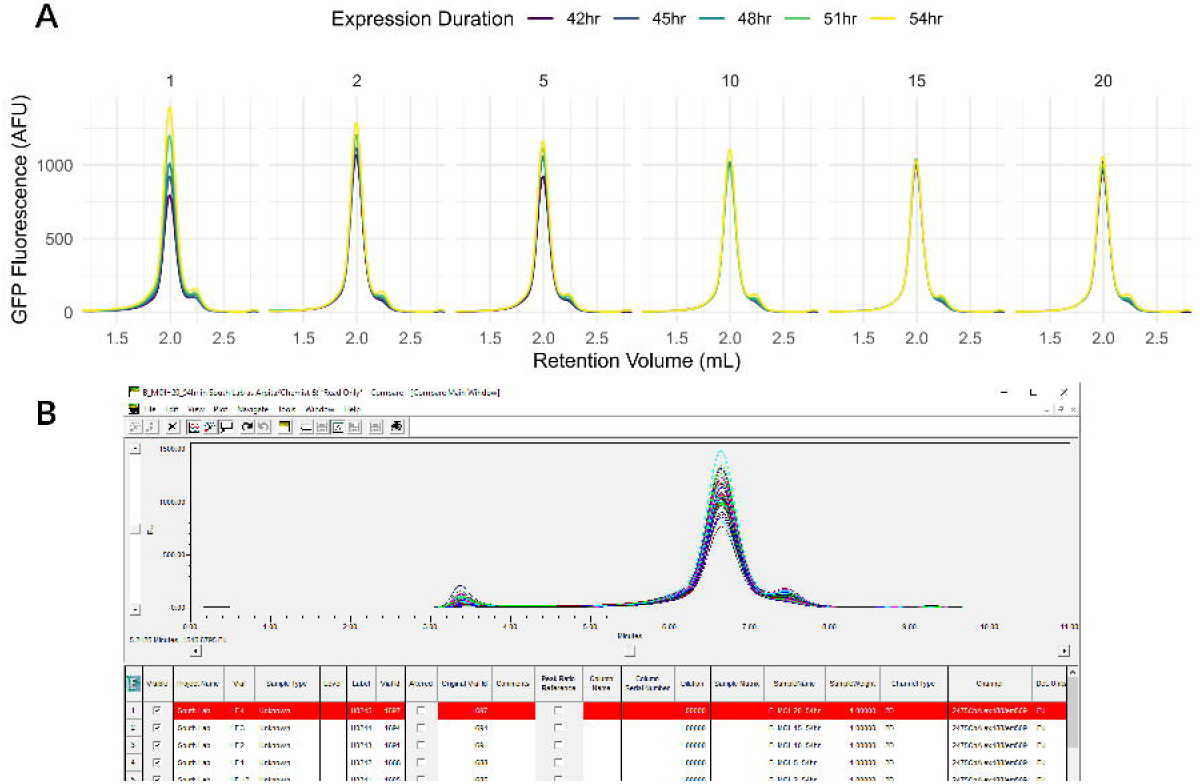
Small cell pellets with the indicated baculovirus MOI and expression duration were thawed, solubilized in DDM, and run in a Waters Acquity Arc Bio HPLC over a Superose 6 Increase 5/150 column. A: Data processed and visualized with Appia and ggplot. Data export and visualization took approximately 20 seconds. Trends in both expression duration and MOI are obvious, and the user can move on with larger-scale expression tests. B: The same data as viewed in the manufacturer software

Manufacturer software, on the other hand, displays all the lines in the same pane (Figure 3B); users must remember which line corresponds to which set of conditions. In this case, it is very difficult to follow the decreasing maximum protein expression with increasing MOI, and even more difficult to notice that the duration effect disappears with higher MOI. For this experiment, we selected an MOI of 2 and 42 hours, even though the maximum is at MOI 1 and 54 hours, as the slight gain from longer expression is not worth the time. This insight was only possible with the clear plot layout enabled by Appia. Additionally, Appia’s standard data format means that even if a similar experiment were run on a separate instrument from a different manufacturer, the same visualization and processing scripts could be used, eliminating the wasted effort of learning a new manufacturer software.

Appia centralizes data analysis workflows. For instance, our lab relies extensively on paired FSEC/SEC assays. First, several constructs of interest are purified in a matrix of conditions. These lysates are assayed for homogeneity and yield by FSEC. All of the resulting plots can be viewed simultaneously, from anywhere with an internet connection, in Appia Web (Figure 4A). We next express a larger quantity of the most promising condition and purify it by affinity chromatography and SEC. Again using Appia Web, we select the most promising fractions of the preparatory SEC (Figure 4B) and assess them again for monodispersity using FSEC (Figure 4C). We can compare the peak profiles directly with previous purifications of the same or different conditions and constructs, or our small-scale tests, by loading up the relevant Experiments in Appia Web. The single-destination simultaneous display of both preparative and analytic data makes comparison and selection both easier and faster, and simultaneous display of raw and normalized data allows for direct comparison of both yield and monodispersity without any need for user input or task switching.

**Figure 4.**
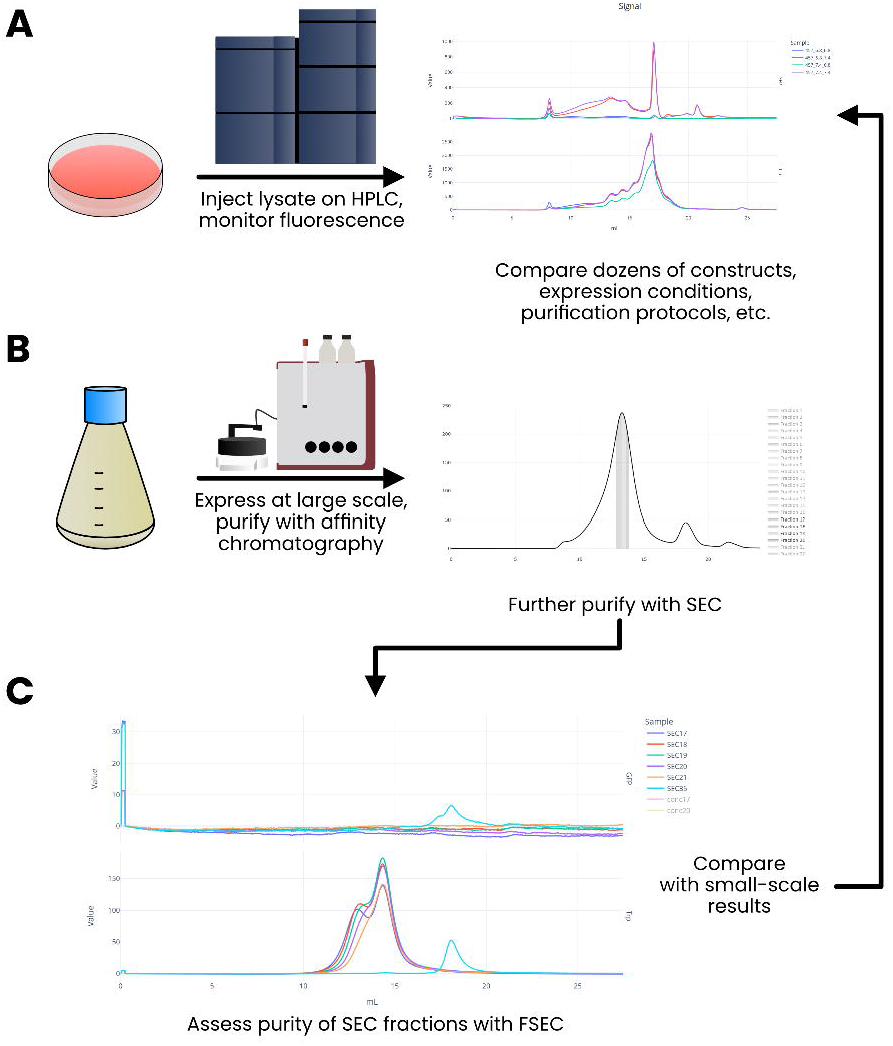
In a paired SEC/FSEC assay, small-scale purifications of varying constructs in varying conditions are analyzed by FSEC for monodispersity and yield (A). Next, conditions of interest are purified by affinity chromatography and preparative SEC (B). Fractions from each step are selected with Appia Web for re-analysis using analytic FSEC (C) to assess the homogeneity of the samples. These results can easily be compared with prior analytic work of other constructs or conditions without ever leaving the Appia Web interface.

Appia Web also makes binding interactions easier to analyze (Figure 5). The user can normalize FSEC traces to the unbound peak. This makes the relative peak heights a proxy for the relative proportion of the fluorescent species in the bound or unbound states. For instance, here we show a binding interaction between the Epithelial Sodium Channel (ENaC) and its regulatory partner NEDD4-2, tagged with GFP (9). ENaC is untagged, so any GFP signal indicates the position of some species containing NEDD4-2. It is plainly clear from the raw traces that there exists some binding interaction between the two, and the traces normalized to unbound peak height give a heuristic approximation for relative binding: at a 1:5 NEDD4-2:ENaC ratio, the unbound peak is about 4 times higher than the bound peak, giving an approximate 20% of the total NEDD4-2 population bound by ENaC.

**Figure 5.**
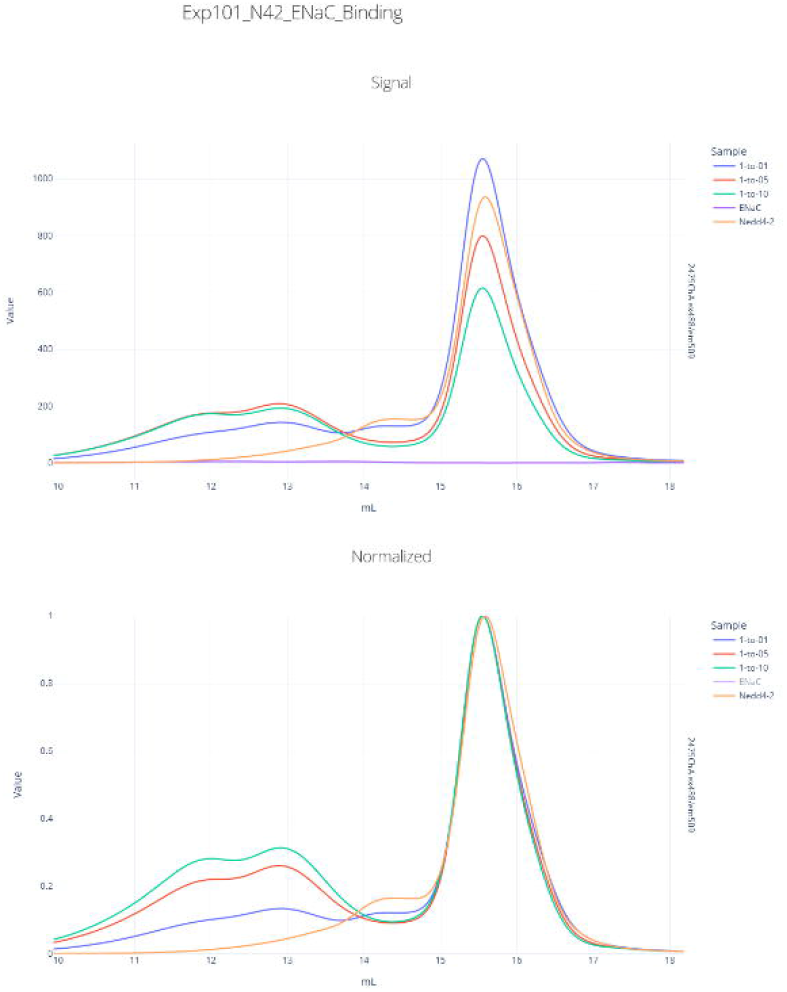
Top: raw GFP fluorescence from NEDD4-2-GFP fusion protein. Bottom: fluorescence signal normalized, setting free NEDD4-2-GFP peak height to 1.

## Discussion

Chromatography is a workhorse assay of chemistry and molecular biology. Because of the assay’s flexibility, manufacturer software is often difficult to use for simpler, routine assays, because it must present functionality required for advanced chromatographic assays. Appia simplifies analysis of experiments where only the simple readouts of peak shape, heigh, and position are required by producing easy-to-read plots, giving users insight into their data with little distraction.

Appia’s standard data formats and web interface also make data sharing very simple. Collaborators can directly view the chromatogram of purified material from their own computer, and download the data for further investigation if needed. Even if two labs use different chromatography instruments, Appia’s standard data format means that there are no extra efforts spent learning a new set of software to analyze the relevant data. Moreover, Appia Web presents simple one-click controls for common data manipulations such as scaling and peak height normalization, meaning that even advanced chromatography users performing standard assays will find Appia easier and faster to use than manufacturer software.

Appia is, of course, not sufficient to run an instrument or process advanced chromatography workflows. Instead, Appia is meant to provide a free, open-source interface to make all parts of day-to-day chromatography simpler: analysis with the streamlined, appealing interface; sharing through Appia Web; and publication through Appia’s plotting functionality and standardized data format.

## Experimental procedures

### Appia

Appia is currently written to analyze data from Waters, Shimadzu, and Agilent HPLCs and GE/Cytiva FPLCs, and is under active development. Support of a new chromatography manufacturer requires writing a parser to convert the manufacturer export format to the Appia format; this typically takes between one and two hours for someone moderately experienced in python, and only has to be done once per manufacturer. Requests for additional manufacturer support are very welcome, and require only submission of some basic information as well as a few representative chromatogram export files. Appia can be installed with a single command. Appia Web is optional, but highly recommended. It requires installation of both Appia and an Apache CouchDB database, along with basic networking. The GitHub repo contains a Docker Compose YML template. Docker Compose automatically builds both the Appia web UI and the CouchDB database and appropriately networks them together with a single command, meaning that installation can be performed with minimal technical expertise. A typically Appia installation might include an installation of Appia on each instrument computer, with some central, web-accessible computer running Appia Web. Each user’s personal computer does not require any additional software to access Appia Web.

### Expression Profiling

HEK293S GnTI^-^ cells were infected with baculovirus for GFP-tagged CMP-sialic acid transporter at multiplicity of infection (MOI) of 1, 2, 5, 10, 15, or 20(10). Cells were incubated at 30 °C for 54 hours, with 2 mL aliquots removed at 42, 45, 48, 51, and 54 hours of infection. Aliquots were centrifuged at 3,000 xg for 10 minutes, after which cell pellets were flash-frozen in liquid nitrogen and held at -80 °C until all pellets were prepared. Cell pellets were solubilized in solubilization buffer (50 mM HEPES pH 7.5, 150 mM NaCl, 0.01 mg/ml deoxyribonuclease I, 0.7 μg/ml pepstatin, 1 μg/ml leupeptin, 1 μg/ml aprotinin, 1 mM benzamidine, 0.5 mM phenylmethylsulfonyl fluoride, and 2% (w/v) n-dodecyl-β-D-maltopyranoside [DDM]) for 1 hour at 4 °C, then centrifuged at 35,000 xg for 30 minutes. Next, 50 μL supernatant was injected on a Superose 6 Increase 5/150 column (Cytiva) on a Waters Acquity Arc Bio HPLC and monitored for GFP fluorescence. Data were processed through Appia, then through a standard plotting script developed previously for this type of assay and provided in the Supporting Information.

### SEC Fraction Analysis

Wild-type-like ENaC δβγ was expressed in HEK293S GnTI^-^ cells and purified as described previously for αβγ ENaC (11). Briefly, cells were infected with baculovirus at an MOI of approximately 2 and incubated at 30 °C for 72 hours, then harvested by centrifugation and flash-frozen. ENaC-GFP cell pellets were solubilized in solubilization buffer (20 mM HEPES pH 7.4, 150 mM NaCl, 2 mM MgCl_2_, 25 U/mL Pierce universal nuclease, protease inhibitors, and 2% digitonin) for 1 hr at 4 °C, then centrifuged at 100,000 xg for 45 minutes. Supernatant was passed over a GFP nanobody resin to bind ENaC-GFP. ENaC was eluted from the resin by treatment with trypsin (33 μg/mL resin) to cleave the channel from GFP. Trypsin-eluted fractions were concentrated and injected on a Superose 6 Increase 10/300 column (Cytiva) on an AKTA Pure FPLC. Fractions were collected and re-analyzed by FSEC on a Waters Acquity Arc Bio. Running buffer was the same for SEC and FSEC: 0.5 mM GDN, 20 mM HEPES pH 7.4, 200 mM NaCl. SEC data and fraction FSEC data were exported and processed with Appia and analyzed entirely in Appia Web.

### NEDD4-2/ENaC Binding

Human NEDD4-2-GFP was expressed in Sf9 cells at an MOI of 5 for 48 hours, then flash frozen. Cell pellets were resuspended in 100 mL TBS (20 mM Tris pH 7.5, 200 mM NaCl) with 25 U/mL Pierce universal nuclease and protease inhibitors per 800 mL cell culture, then sonicated as follows: 5 minutes total sonication, 10 seconds on, 30 seconds off, power level 7. The lysate was centrifuged at 45,000 xg for 45 minutes, then bound to 10 mL TALON resin per 800 mL cell culture. The resin was washed with 5 CV TBS, then 2 CV TBS + 10 mM imidazole, and then NEDD4-2-GFP was eluted with TBS + 250 mM imidazole. NEDD4-2-GFP was concentrated and further purified by SEC. ENaC αβγ was purified as described above for SEC Fraction Analysis.

NEDD4-2-GFP and ENaC were mixed at varying ratios and analyzed by FSEC on a Waters Acquity Arc Bio HPLC in TBS + 0.5 mM DDM. Final NEDD4-2-GFP concentration was held constant at 71.2 nM, while ENaC was added in equal amounts, five-fold excess, or ten-fold excess, then 50 μL of these mixtures were injected onto a Superose 6 Increase 10/300 column (Cytiva). The GFP channel was monitored to detect a shift in NEDD4-2-GFP elution position. The earlier-eluting peak that increases in height and area with increasing ENaC concentration elutes before both the ENaC alone and NEDD4-2-GFP alone peak positions in tryptophan fluorescence (not shown), indicating that the species is larger than either protein on its own.

Data were exported from Waters Empower and processed by Appia, then analyzed entirely in Appia Web.

## Data availability

Raw chromatography data are available upon request to IB. Appia source code is available at <u>DOI: 10.5281/zenodo.6975032</u> under the MIT license. Appia is written in the following languages and packages under their respective licenses: python, PSF license; R, GPL-2 | GPL-3; tidyverse, MIT; Plotly, MIT. Copies of the relevant licenses are available in the Appia GitHub repo.

This article contains supporting information.

## Acknowledgements

We are extremely thankful to the OHSU Medicinal Chemistry Core for providing sample Agilent data during development, and for helpful comments and discussion. Thanks also to Eric Gouaux, Steve Mansoor, and Shivani Ahuja for Shimadzu data. We would also like to thank the lively community of python, plotly, R, and ggplot users. Without the wealth of knowledge and help provided by these communities, Appia could never have been written. This paper would lose valuable biological perspective without the example chromatography data kindly provided by James Cahill and Alex Houser. Finally, we would like to thank Alex Houser, James Cahill, Kim Hartfield, Sigrid Noreng, Arpita Bharadwaj, for valuable feature suggestions and frequent bug reports. This work was supported by the National Institute of General Medical Sciences (T32GM071338, initially to support RP, and R01GM138862 to IB). The content is solely the responsibility of the authors and does not necessarily represent the official views of the National Institutes of Health.

The authors declare that they have no conflicts of interest with the contents of this article.

## Footnotes

SEC: Size-Exclusion Chromatography
FSEC: Fluorescence Size-Exclusion Chromatography
CSV: Comma Separated Value
MOI: Multiplicity Of Infection
ENaC: Epithelial Sodium Channel
NEDD4-2: Neuronal precursor cell-expressed developmentally downregulated 4-2; an E3 ubiquitin ligase
HPLC: High-Performance Liquid Chromatogram, typically used for analytic chromatography
FPLC: Fast-Performance Liquid Chromatogram, typically used for preparative chromatography

